# Emerging phylogenetic structure of the SARS-CoV-2 pandemic

**DOI:** 10.1101/2020.05.19.103846

**Authors:** Nicholas M. Fountain-Jones, Raima Carol Appaw, Scott Carver, Xavier Didelot, Erik Volz, Michael Charleston

## Abstract

Since spilling over into humans, SARS-CoV-2 has rapidly spread across the globe, accumulating significant genetic diversity. The structure of this genetic diversity, and whether it reveals epidemiological insights, are fundamental questions for understanding the evolutionary trajectory of this virus. Here we use a recently developed phylodynamic approach to uncover phylogenetic structures underlying the SARS-CoV-2 pandemic. We find support for three SARS-CoV-2 lineages co-circulating, each with significantly different demographic dynamics concordant with known epidemiological factors. For example, Lineage C emerged in Europe with a high growth rate in late February, just prior to the exponential increase in cases in several European countries. Mutations that characterize Lineage C in particular are non-synonymous and occur in functionally important gene regions responsible for viral replication and cell entry. Even though Lineages A and B had distinct demographic patterns, they were much more difficult to distinguish. Continuous application of phylogenetic approaches to track the evolutionary epidemiology of SARS-CoV-2 lineages will be increasingly important to validate the efficacy of control efforts and monitor significant evolutionary events in the future.

## Introduction

The rapid spread of the novel coronavirus SARS-CoV-2 since December 2019 represents an unparalleled global health threat^1^. Within four months of emerging from Wuhan in Central China, SARS-CoV-2 has now spread to nearly every country and is a major source of mortality^1^. The first cases of the virus outside China occurred in Thailand on January 13, and by January 30 there were 83 cases in 18 countries. As of May 19, there were over 4.5 million cases in 203 countries or territories^1^. Coronaviruses (order: Nidovirales, family: Coronaviridae) are enveloped positive-sense non-segmented RNA viruses that infect a variety of mammals and birds. SARS-CoV-2 is the seventh coronavirus to be identified infecting humans. The closest relatives (RaTG13 and RmYN02, 96% and 93% nucleotide identity respectively) derive from the Intermediate Horseshoe bat (*Rhinolophus affinis*) and the Malayan Horseshoe bat (*Rhinolophus malayanus*)^2^, although the original host is yet to be conclusively identified ^3^. Since spilling over to humans, the virus has diverged rapidly, but it is unclear whether these mutations have resulted in SARS-CoV-2 lineages with different epidemiological and evolutionary characteristics ^4–9^. Several lineages have been highlighted for potential significance ^4–6,9^. For consistency, we adopt the nomenclature outlined in ^8^ which classifies the initial lineages as A and B (previously labelled ‘S’ and ‘L’^4^). There is some evidence that Lineage A is ancestral to the more recent Lineages B ^8^, even though the earliest assembled genomes from December 2019 belong to Lineage B ^4,8^. Sequences within Lineage A and the closest known bat virus share two nucleotides in ORF1ab and ORF8 genes that are not found in Lineage B ^8^. More recently, a new lineage ‘G’^6^ has been documented originating Europe in February ^6^. For consistency we call this Lineage C. It is currently unclear if these lineages differ phenotypically, or whether these lineages show distinctive demographic signatures (i.e., diversity increasing, plateauing or declining). Any further population sub-structure within these three lineages is also unknown at this point.

Pathogen population structure can provide key insights into the epidemiology of an outbreak, such as whether intervention strategies are working to contain spread ^7^. Population structure may also align with geography, reflecting the contact structure of the host population. Understanding these variations is important both for vaccine development and evaluating the impact of control efforts across the globe. Detecting structure, particularly in recently emerged outbreaks, is a challenge as these patterns within the data can be cryptic ^8^. For example, some lineages within a population can be rapidly expanding whereas others can be stationary ^8^. Utilizing large numbers of sequences provided by GISAID ^9^and recently developed phylodynamic tools, we interrogate SARS-CoV-2 population patterns to identify ‘hidden’ structure in the pandemic and investigate whether lineages are geographically partitioned and/or are on distinct demographic trajectories.

### Three distinct lineages

Our analyses show support for three distinct lineages of SARS-CoV-2 actively spreading around the world (Fig. 1). We demonstrate that they are not only phylogenetically separate but also have different demographic trajectories. Based on our maximum likelihood and Bayesian time-scaled phylogenies (see *Methods*), we estimated that Lineage A (and SARS-CoV-2 overall) diverged from its most recent common ancestor (MRCA) in November 2019 (95% high posterior density/confidence intervals November – December 2019, Fig. 1). Estimates from both approaches are comparable to other studies that have analysed greater numbers of sequences ^9^. Since emerging in China, our demographic analysis ^13^ suggests that the growth rate of the effective population size of Lineage A increased in early January (Fig. 2a), then decreasing throughout February before increasing once more. This dip coincides with control of the pathogen in China ^14^ and subsequent uncontrolled spread in Europe and North America. We found a similar pattern when we analyzed the complete dataset (Fig. 3a). The majority of sequences belonging to Lineage A originated from China in January to early February, whereas sequences from the US, and Washington State in particular, make up the majority of the sequences collated in March.

**Fig. 1.**
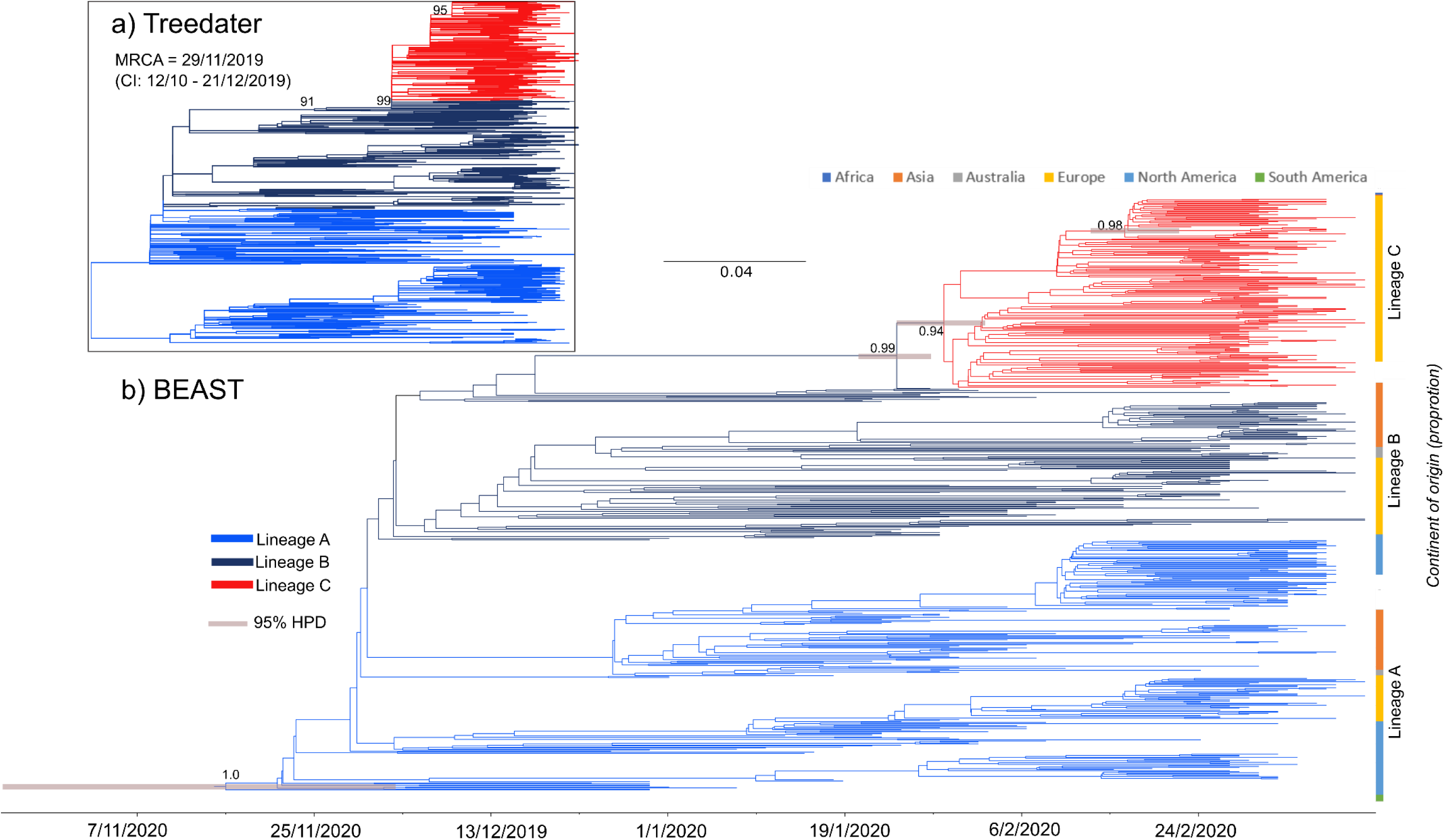
*Treedater* maximum likelihood tree (a) and Bayesian time-scale phylogeny (b) revealing the three SARS-CoV-2 lineages we identified with unique demographic signatures (Lineages A, B & C). Branches in both trees are coloured by lineage identified using the *treedater* phylogeny (see *Methods* for details). Most recent common ancestor (MRCA) estimates from the *treedater* analysis are also provided. Density bars are shown representing the 95% highest posterior density (HPD) intervals for the dating of each lineage. Node posterior support values and bootstrap support values are shown for internal nodes not leading to leaves with values > 0.8 or 80% posterior or bootstrap support respectively. See Fig. S1 for the Bayesian tree with all posterior support values. Stacked bar plots show the proportion of sequences from each country classified in each lineage.

**Fig. 2.**
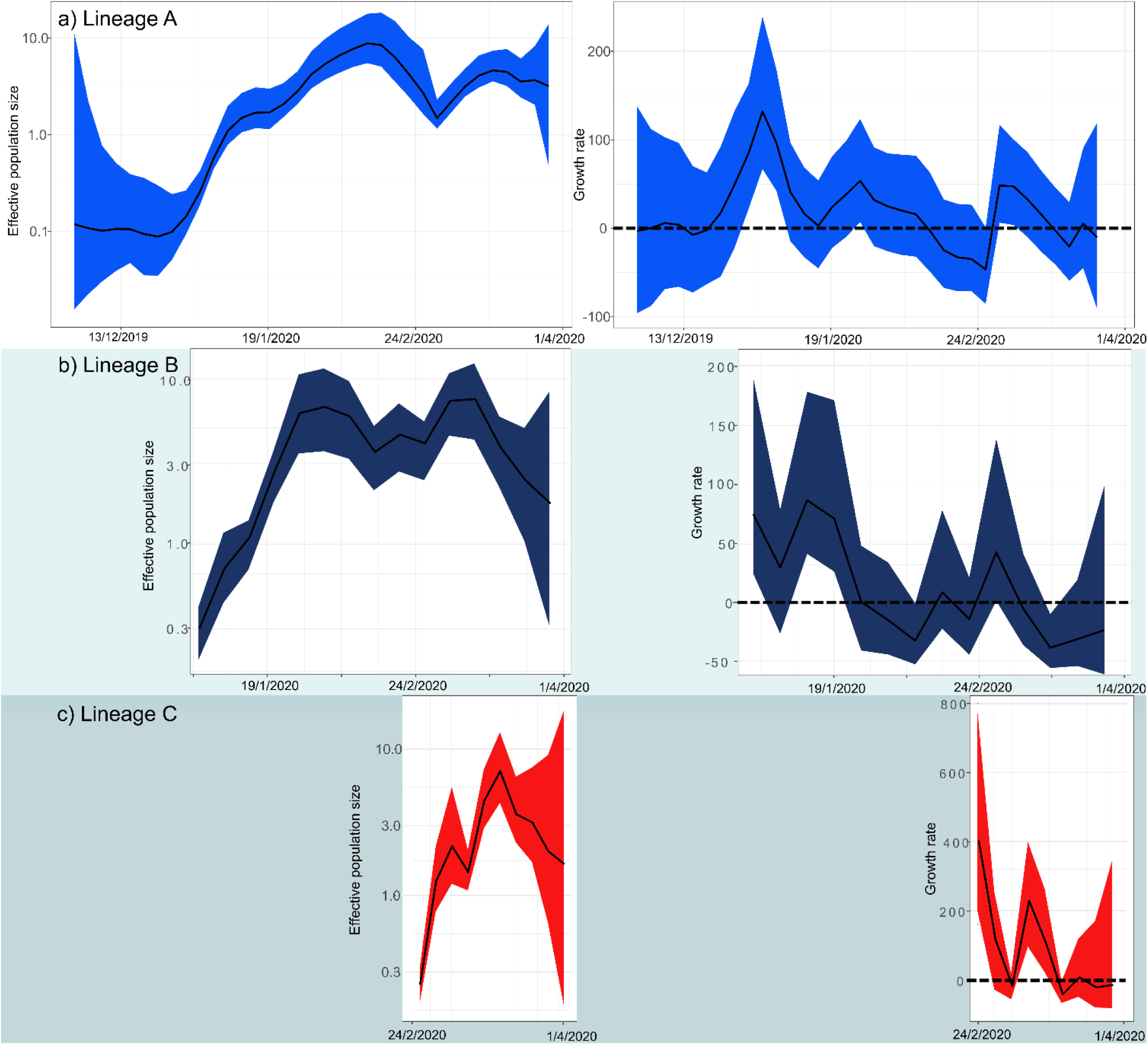
Effective population size (left panels) and growth rate of the effective population size per year (right panels) estimated through time for the three identified SARS-CoV-2 Lineages from our *skygrowth* models. The coloured 95% high probability density (HPD) intervals reflects lineages identified in Fig. 1. Dashed lines in the left panels indicate a growth rate of zero.

**Fig. 3.**
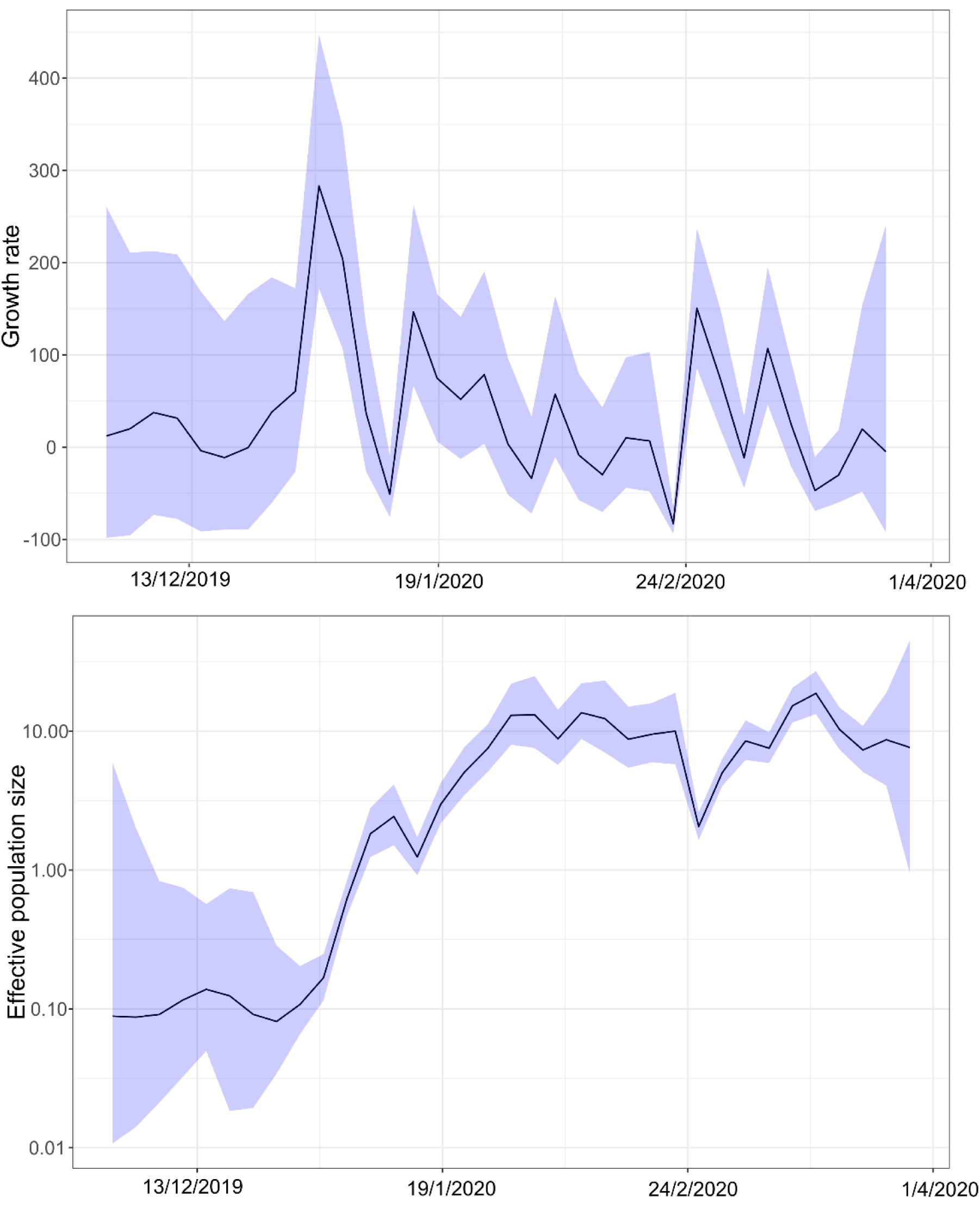
Growth rate (a) and effective population size (b) estimates through time from our *skygrowth* model using the complete dataset (all lineages of SARS-CoV-2). Light blue shading represents the 95% HPD of the estimates.

Our results support other analyses suggesting that Lineage B was derived from Lineage A and was not an independent introduction, even though Lineage B contains the earliest available genomes ^4,8^. Linked mutations in ORF1ab (8782, synonymous) and ORF8 (28144, non-synonymous) help to separate these lineages (as in ^4,8^). Non-synonymous mutations in ORF14 (28881-3) also partially define these lineages. However, there is a high degree of phylogenetic uncertainty about the node representing their most recent common ancestor and this lineage may be polyphyletic (Fig. 1, Fig. S1). Soon after diverging from Lineage A, the growth rate of Lineage B was at its highest but then formed a pattern of peaks and troughs with the credible interval including zero (representing no growth) from January onwards (Fig. 2b). The peak growth rate coincided with that of Lineage A (Fig. 2) indicating that this first wave of SARS-CoV-2 through China generated a relatively large amount of the genetic diversity. As many sequences classified in Lineage B originate from China (Fig. 1) the subsequent decline of this lineage may also be linked to control of the virus. There is also evidence for a spike in growth rate of both Lineage A and Lineage B when spread increased outside of China and this coincides with the divergence date for Lineage C (Fig. 2).

Lineage C, in contrast, was predominantly European with no evidence that it circulated in China (Fig. 1). This lineage was well supported as monophyletic (node posterior support = 0.99, 91% bootstrap support, Fig. 1) and diverged from Lineage B in late January (95% HPD late January – early February). Linked non-synonymous mutations differentiated this lineage in the S gene (sites 23402-04) and ORF1ab (14407-09) regions. The mutations in the ORF1ab gene alter the RNA-dependent RNA polymerases (RdRp) that are crucial for the replication of RNA from the RNA template. There is evidence that this RdRp mutation may increase the mutation rate of the virus overall by reducing copy fidelity^7^. The growth rate of Lineage C was initially high in late February, prior to the rapid increase of cases in Europe, but then declined, with one further peak around February 27, although the short duration suggests this may not be significant and could represent sampling noise. Accordingly, the effective population size of Lineage C increased rapidly during February – March, whereas there was only a small increase estimated for Lineage A and a decline in Lineage B (Fig. 2). Real-time phylogenetic reconstruction in Nextstrain ^15^ has subsequently shown (as of May 18 2020) that this lineage has further expanded and is the most frequently sampled across the globe. Given the potential significance of the RdRp and spike gene mutation and subsequent rapid expansion across the globe, our results are consistent with the hypothesis that mutations in Lineage C have increased transmissibility compared to the other lineages ^6^. However, our results on their own can not rule out the possibility that the expansion of this lineage was the result of a founder effect ^6^ (i.e., the rapid expansion of Lineage C was because it was the first broadly transmitted in Europe). More analysis of these co-circulating lineages at a local level would be required to evaluate the relative fitness of the lineages.

### The growth and decline of SARS-CoV-2 lineages

We were able to identify three lineages that were not only genetically distinctive but also had unique demographic signatures, revealing insights into the underlying epidemiology of this pandemic. The number of cases increases day-by-day, as does the effective population size of the virus overall (Fig. 3b); both to be expected by their linear relationship in the early phase of a susceptible-infected-removed (SIR) compartmental model ^16^. It appears that this increase is not distributed evenly across the phylogeny, with all lineages showing some evidence of decline at different times. However, there is bias in countries represented in the GISAID dataset we accessed, with, for example, no sequences in our dataset from the Middle East even though there was a significant (and ongoing) outbreak in this region. Even though the outbreak is only months old at the time of writing, there is already sufficient genetic diversity to track the demographic trajectories of each lineage. Approaches such as the one presented here, combined with workflows quantifying geographical lineage dispersal ^17^, will be even more useful in the coming months to assess the longer-term impacts on SARS-CoV-2 control measures across the globe.

## Methods

We downloaded 779 complete “high coverage only” SARS-CoV-2 genome sequences from GISAID ^9^ (Global Initiative on Sharing All Influenza Data; https://www.gisaid.org/) on the 24^th^ March 2020. We aligned these sequences with MAFFT ^18^ using the CIPRES ^19^ server and visually checked the results. We trimmed the first 130 bp and last 50 bp of the aligned sequences to remove potential sequencing artefacts in line with Nextstrain protocol ^15^. We tested for recombination in our alignment using RDP4^120^. We removed all duplicate sequences and sequences with more than 10% missing data. We then constructed a Maximum Likelihood tree using IQ tree with 1000 ultrafast bootstraps^21^ using the inbuilt model selection algorithm (‘ModelFinder’ ^22^). We confirmed that there was a significant temporal signal in the dataset using root to tip regressions in TempEst^21^ (R^2^ = 0.19, correlation coefficient = 0.42). We removed sequences from Washington State and China that likely had some sequence error as they were strong outliers in the TempEst analysis. This reduced the dataset to 587 complete SARS-CoV-2 genomes.

We used both the maximum likelihood-based *treedater* method ^24^ and a Bayesian approach to reconstruct the timing and spread of SARS-CoV-2. We employed the computationally intensive Bayesian methodology (BEAST version 1.10.4 ^25^ with BEAGLE ^26^ computational enhancement) to validate our maximum likelihood MRCA estimates and to provide dating estimates for internal nodes of interest. For the BEAST analysis, as there is strong evidence that the pandemic is growing, we assumed an exponential growth coalescent model. Preliminary analyses revealed that there was no statistically significant difference in evolutionary rate estimates using either a strict or relaxed molecular clock. Thus, we assumed a strict molecular clock and used uninformative continuous-time Markov chain (CTMC) reference prior to estimate substitution rate across time. We also tested a relaxed clock model and found very little rate variation across the tree and very similar estimates of the MRCA. We performed each BEAST analysis in duplicate and ran the MCMC chains for 200 million iterations sampling every 20 000 steps. We visualized these results using Tracer ^27^ and ensured that all parameter estimates had converged with an effective sample size (ESS) > 200. We generated a maximum clade credibility (MCC) tree using TreeAnnotator, discarding 20% as burn-in.

Our previously described ML tree was used as input of the *treedater* method ^24^ to produce a ML time-scaled phylogeny. *Treedater* is an efficient maximum likelihood method that implements both a strict clock model using a Poisson process and a relaxed clock model using a Gamma-Poisson mixture. We estimated the confidence intervals for the dates of ancestors in this tree using parametric bootstraps.

We then used this time-stamped ML tree to test for structure within the tree using the non-parametric *treestructure* approach ^11^. Briefly, this method partitions the tips and internal nodes of a tree into discrete sets characterized by comparable coalescent patterns. See ^11^ for analytical details. Given the relatively low levels of genetic diversity, we constrained our structure analysis to be able to identify a maximum of four lineages by making the minimum clade size 145 sequences and performed 100,000 tree simulations (with a significance threshold of 0.05). We could not perform the same analysis on our Bayesian tree as it included negative branch lengths, which violates the assumptions of *treestructure*.

For the complete dataset and each lineage subset, we modelled the effective population size growth rate through time using the *skygrowth* package^11^. *Skygrowth* is a non-parametric Bayesian approach that applies a first-order autoregressive stochastic process on the growth rate of the effective population size. We parameterized our *skygrowth* models assuming that SARS-CoV-2 effective population size could change every three days. We used an exponential distribution with a mean of 0.1 to estimate the precision parameter (Tau). We ran the MCMC for 20 million generations thinning every 1000^th^ sample and considered each analysis to be converged if the ESS >200. We compared our *skygrowth* models to *Skygrid* models using the R package ‘phylodyn’ ^24^ using the default settings. The ML tree and code used to perform these analyses are available here: https://github.com/nfj1380/covid19_evolution.

## Acknowledgements

We thank all the laboratories that provided sequences to the GIAID project. This project was supported by an Australian Research Council Discovery Project Grant (DP190102020).

