## Supplementary material for "Emerging phylogenetic structure of the SARS-CoV-2 pandemic": Fig. S1, Fig. S2

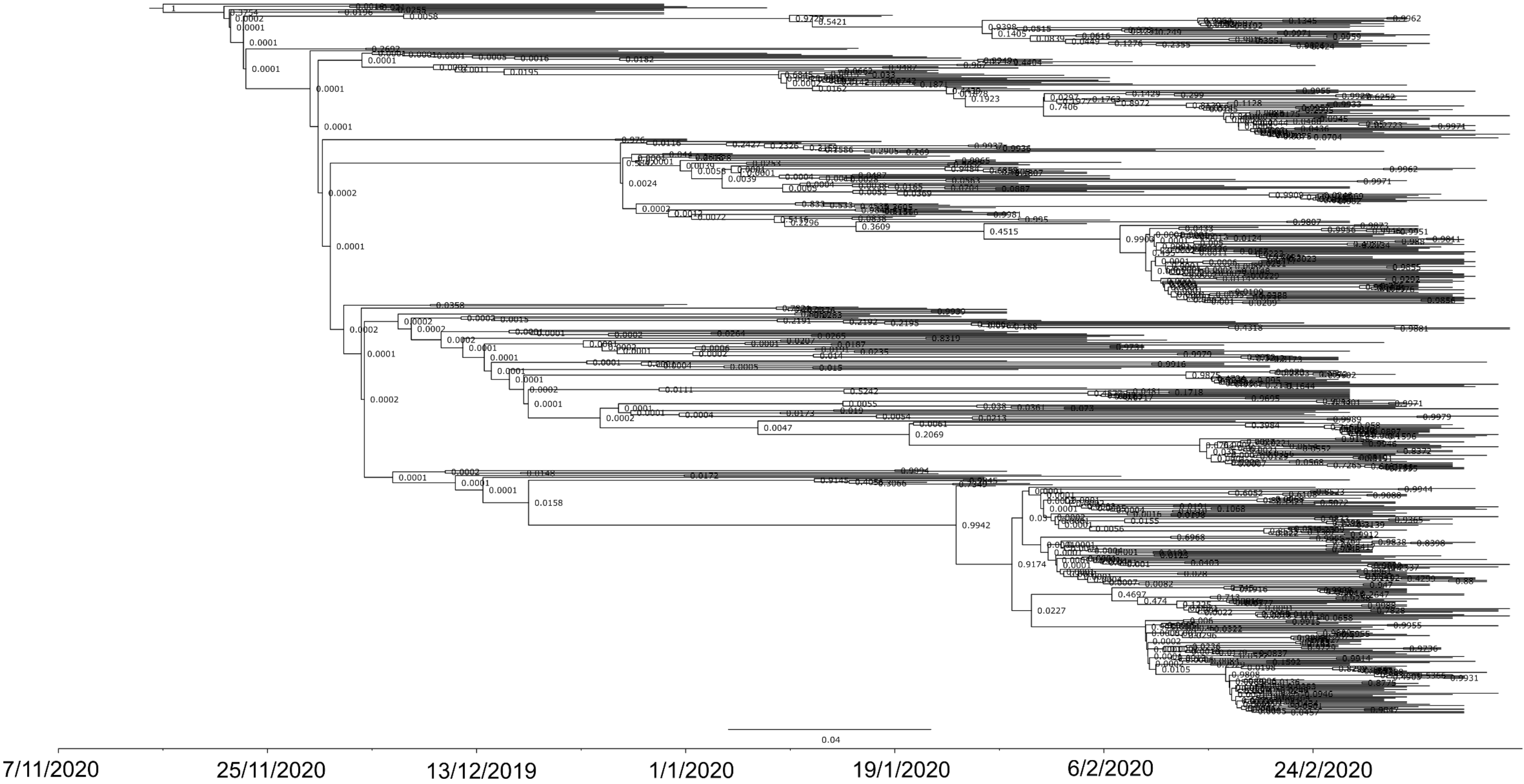

**Fig. S1.** Time-scaled Bayesian phylogeny showing all posterior support values.

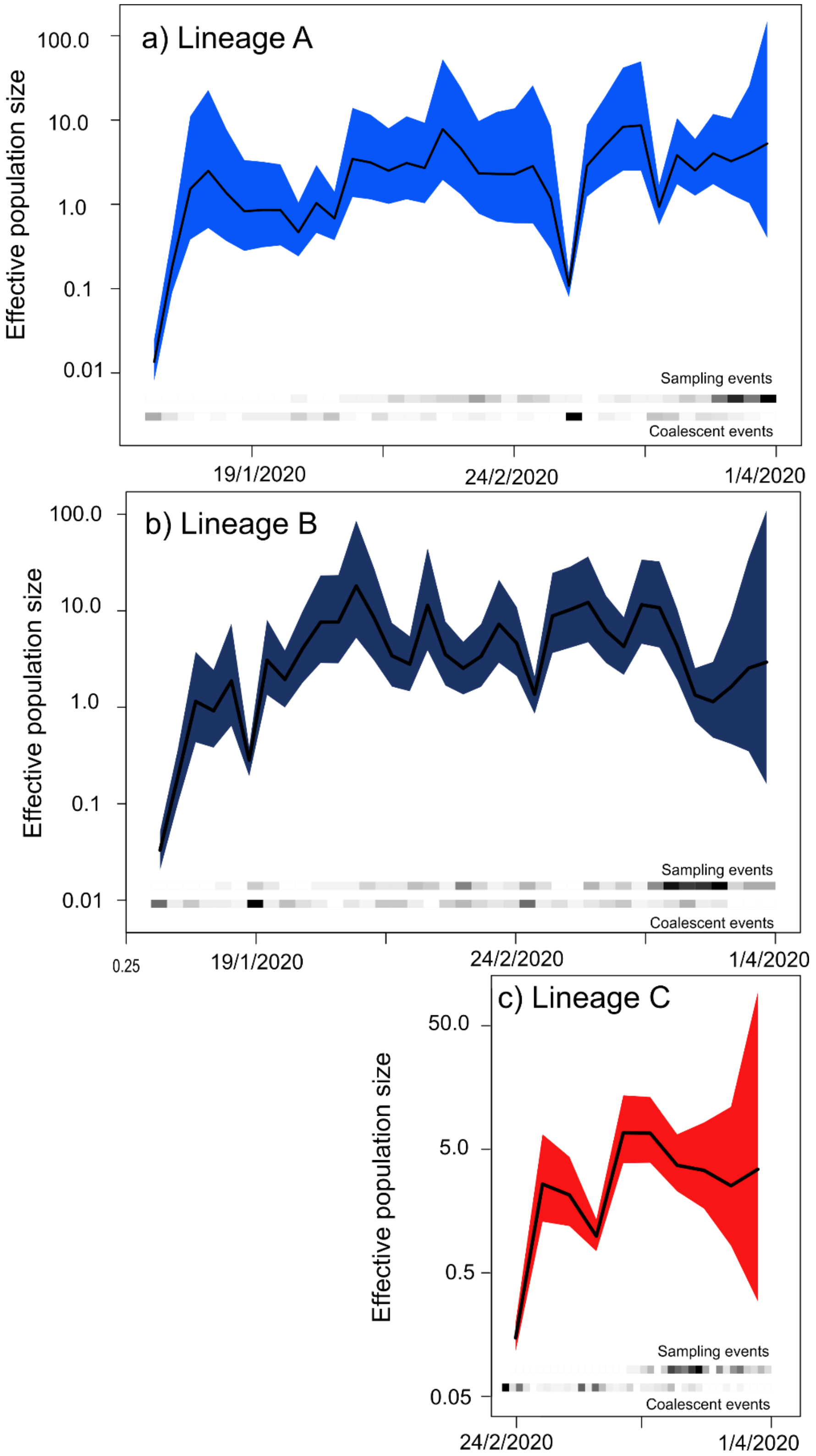


**Fig. S2.** Skygrid plots of the three identified SARS-CoV-2 lineages showing effective population size through time using the *phylodyn* approach [1]. The coloured 95% high probability density (HPD) intervals reflect lineages identified in Fig. 1.
